# Empirical modelling of trait selection by partitioning selection into direct selection and selection that is mediated by interspecific interactions

**DOI:** 10.1101/045583

**Authors:** Christian Damgaard

## Abstract

Trait selection has received considerable attention in the pursuit to understand niche-based community assembly processes and to generate ecological predictions. To further advance the study of trait selection, a conceptual statistical model is presented that outlines and discuss the possibilities of i) estimating the effect of interspecific interactions on traits rather than just testing weather selection has had an effect on the observed trait distributions, ii) discriminating between environmental filtering and niche partitioning processes and estimate the characteristic features and importance of both processes, and iii) predicting the effect of environmental changes and gradients on trait selection. To achieve these goals a number of necessary assumptions have to be specified and these assumptions are discussed and assessed. Simulated plant cover data from a simple uniform environment was successfully fitted to the model and the results indicates that it is possible to partition direct population growth and population growth that is mediated by interspecific interaction. The data requirements of the model are modest, i.e. time series data on plant species abundance and a species trait matrix. Consequently, the model concept may be used to model trait selection, including the effect of interspecific interactions, in many existing plant ecological datasets.

## Introduction

Interspecific interactions among neighbouring plants typically arise because the resources needed for plant growth and reproduction are limited, and the plant that extracts or monopolizes most of the limiting resources will grow faster and reproduce in greater numbers (e.g., Goldberg et al., 1990; Weiner, 1986).

The possibly important role of interspecific interactions in regulating natural plant communities and determining community assembly rules (e.g., Gotelli and McCabe, 2002; Kraft et al., 2015b; Silvertown et al., 1999; Weiher et al., 1998) has been investigated in a multitude of studies using different methods (Damgaard, 2011). However, considering its high ecological relevance and status as a classic research question in plant population ecology, it is noteworthy that only relatively few studies have measured the direct effect of interspecific interactions on plant performance and its role for regulating plant communities in undisturbed natural communities, and the results are still too sparse to allow much generalization across different plant communities or even among years (Turnbull et al., 2004). This paradox is due to the fact that the measurement of interspecific interactions in natural ecosystems is a non-trivial task (Damgaard, 2011), and applicable methods for measuring interspecific interactions in natural ecosystems is needed in order to make progress in understanding community assembly rules and making quantitative ecological predictions on the effect of environmental changes on biodiversity.

An increasingly popular way of describing plant communities is to focus on the expressed phenotypes of the plant species, i.e. plant traits, rather than on the species itself. The advantage is that plant traits are characteristic features, which to a certain extent will determine the survival, growth and reproductive strategies of the species, and are expected to respond in a more predictable way to an altered environment than the observed change in species composition (Garnier et al., 2004; Garnier et al., 2016; Shipley, 2010a). Furthermore, plant traits involved in resource acquisition and use at the species level will scale-up to ecosystem functioning, provided that traits are weighed by the species’ contribution to the community (Garnier et al., 2007; Lavorel and Garnier, 2002).

Broadly speaking, a trait selection response is caused by i) environmental or biotic filtering processes where the abiotic and biotic environment selects for a certain combination of plant traits that have a relatively high adaptive value in the specific environment independent of the other plant species in the population, i.e. the fundamental niche (Hutchinson, 1957), and ii) competitive or facilitative processes where the trait selection response depends on the traits of the other plant species in the population, i.e. the realized niche (Hutchinson, 1957). The resulting observed selection response on individual traits after both selection processes has operated may be classified into either i) directional selection, where either relatively high or low trait values are favored, ii) stabilizing selection, where specific intermediary trait values are favored over all other trait values, or iii) disruptive selection, where extreme values for a trait are favored over intermediate values.

It is important not to confuse the selection processes with the resulting observed selection response, since multiple assembly processes has been shown to lead to the same pattern of trait dispersion and the same process can lead to different patterns of trait dispersion (Herben and Goldberg, 2014). However, if would be valuable to be able to distinguish between the two types of selection processes from observed changes in the distribution of plant traits since the two different selection processes lead to different expectations of community dynamics including species coexistence and niche-based community assembly processes (e.g. Chesson, 2000; Mayfield and Levine, 2010).

The trait selection process has previously been described by a two-step process in a meta-community model, where plants from a regional species pool are dispersed to a local habitat, and trait filtering excludes individuals with unfit trait values, and within the local species pool, trait values may influence performance, which may lead to patterns of trait convergence or divergence (e.g. Bernard-Verdier et al., 2012; Webb et al., 2010). The selection due to performance differences in the local species pool is thought to be mediated by interspecific interactions as the difference between the fundamental niche and the realized niche of the local species. Under this framework, the effect of interspecific interactions is detected from deviations of the observed trait distribution from random expectations in the local species pool. If the variance of the observed trait distribution is lower than the random expectations, this is an indication of directional or stabilizing selection (convergent trait distribution pattern). Conversely, if the variance of the observed trait distribution is higher than the random expectations, this is an indication of disruptive selection (divergent trait distribution pattern).

Using such test procedures, several plant ecological studies have reported non-random trait dispersion distributions in favor of different niche-based community assembly hypotheses compared to the neutral hypothesis of plant community assembly (Weiher et al., 2011). However, this test procedure has been criticized by e.g. Adler et al. (2013), who argue that trait dispersion tests have low power to detect niche partitioning, and that patterns typically interpreted as either environmental filtering or niche partitioning may be generated by the same process. Most importantly, Adler et al. (2013) note that: “The common interpretation is that species interactions play no role in the abiotic environmental filtering process, while abiotic factors play no role in the competitively driven niche partitioning process. However, the dichotomy between environmental filtering and niche partitioning can arise from an arbitrary decision about the spatial scale of analysis, not from distinct biological processes”.

In a seminal work using maximum entropy models Shipley (2010a; 2010b) estimated the selection response from change in plant abundance. The maximum entropy models have the large advantage that it is not necessary to specify detailed models on selection mechanisms or how the different traits interact (Baastrup-Spohr et al., 2015; Shipley, 2010a; Shipley, 2010b), but this advantage is also their main drawback, since the method does not allow for discriminating between different selection models or whether selection is occurring due to environmental filtering or niche partitioning processes.

Consequently, in order to make progress in the understanding of the role and nature of niche-based community assembly processes in the structuring of plant communities, it would be beneficial to be able i) to estimate the effect of interspecific interactions on traits rather than just testing whether selection has had an effect on the observed trait distributions, ii) to discriminate between environmental filtering and niche partitioning processes and estimate the characteristic features and importance of both processes, and iii) to predict the effect of environmental changes and gradients on trait selection.

To meet these objectives, I present a method for estimating the effect of species trait values on observed population growth in a plant community by estimating parameters in two complementary population growth functions, which partition the observed change in trait distribution of plant population into i) a direct selection process that is independent of the trait distribution of the plant population, which mainly is assumed to arise from environmental filtering processes, and ii) a selection process mediated by interspecific interaction that depend on the trait distribution of the plant population, which mainly is assumed to arise from the niche partitioning processes of competition and facilitation. The resulting model is a one-step trait selection process where the effects of plant traits on population growth is estimated from simple longitudinal plant cover data in an approach that is similar to the approach suggested by Lande and Arnold (1983) to measure selection on correlated characters, but where the effect of traits on population growth is partitioned into direct population growth and population growth that is mediated by interspecific interaction (also see Laughlin et al., 2015; Laughlin et al., 2012). The model operates locally and is conceptually simpler than the two-step meta-community model that previously has been used (e.g. Bernard-Verdier et al., 2012; Webb et al., 2010). Furthermore, a one-step trait selection response is probably a more realistic model of the selection process, since there are no compelling reasons for why the processes of environmental filtering and niche partitioning should not operate simultaneously.

The modest aim of this paper is only to present the model concept and for demonstration purposes to apply it on a toy example. As explained later there are a multitude of possible selection processes, that may be modelled using the model concept and it is meaningless to explore the fitting properties of all the different combinations; except in the context of a genuine plant ecological example.

## Model

A plant community has *n* plant species that are characterized by *m* species-specific plant traits, which are known to be important for plant growth and demography. The plant traits are stored in a species-trait matrix, ***T**_n,m_*, with *n* rows and *m* columns.

The relative local abundance of the plant species is measured by either biomass or cover at time *t, q_j,t_*, where 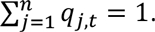. The plants grow, die and reproduce under the influence of interspecific interactions at a given environment where certain combinations of plant traits have a positive effect on growth and reproduction, and other combinations of plant traits have a negative effect on growth and reproduction (compare with Lande and Arnold, 1983).

**Fig. 1.**
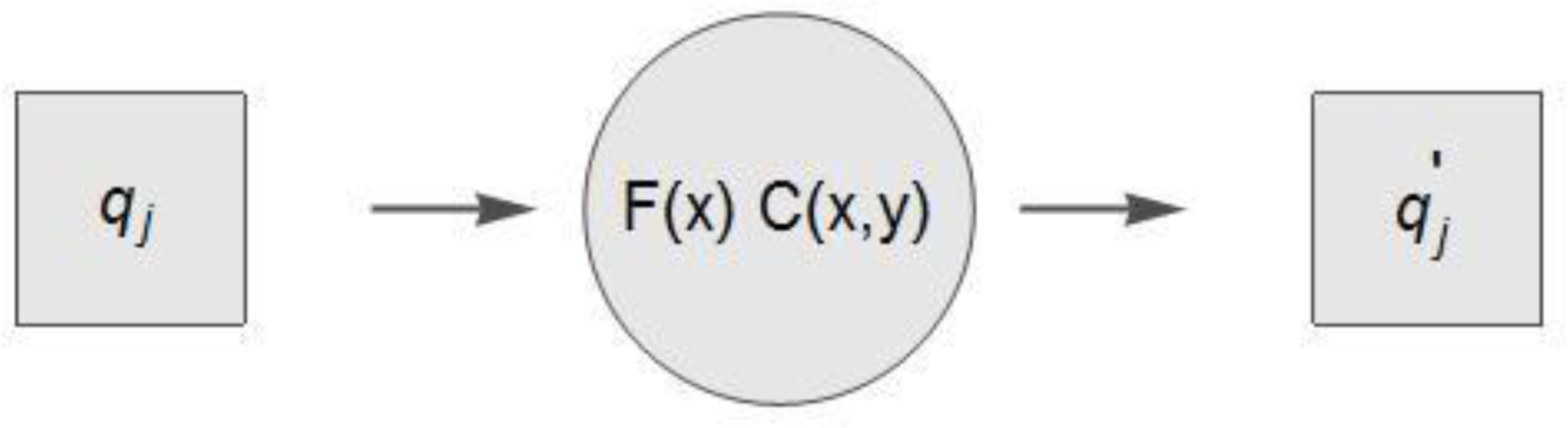
Conceptual figure of the one-step selection model, where *q_j_* is the cover of plant species *j* with trait values *t_k_ = x, q’_j_* is the predicted cover of plant species *j* the following year under the influence of both direct selection forces, *F*(*x*), and selection forces mediated by interspecific interactions, *C*(*x*,*y*).

The predicted cover the following year of plant species *j* with trait values *t_k_ = x* is determined by (Fig. 1),

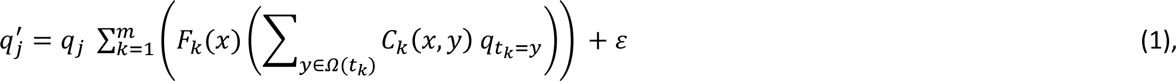

Where *F_k_*(*x*) is the direct population growth function of plant species with trait value *x* for plant trait *k,* and *C_k_*(*x,y*) is a function that models the effect of interspecific interactions on population growth of plant species with trait value *x* for plant trait *k,* where the interspecific interaction of plant species with trait value *x* and *y* is modelled by a distance function, i. e. the effect of species interaction between two species on population growth is determined by the difference in trait values between the two species.

The population growth functions *F_k_*(*x*) and *C_k_*(*x,y*) may vary according to plant life forms, habitat type, and existing prior knowledge of e.g. the type of selection on the different traits. For example, if there is prior information that suggest that directional selection may be operating then this model may be relevant, 
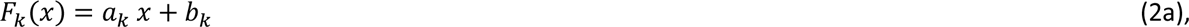

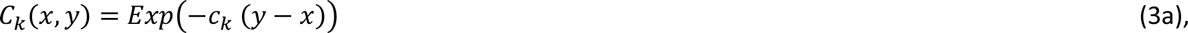
and the following model may be relevant in the case of stabilizing selection, 
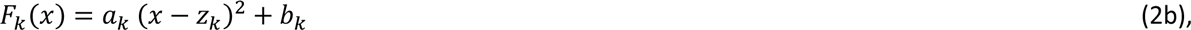

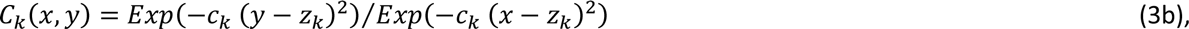
where *z_k_* is an optimum intermediary trait value. Likewise the following model may be relevant in the case of disruptive selection, 
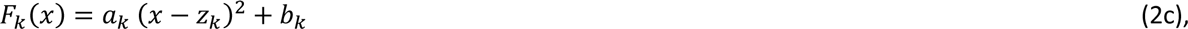

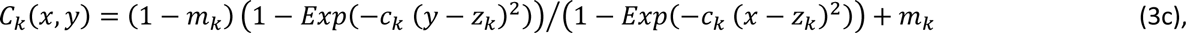
where *z_k_* is a minimum intermediary trait value with fitness *m_k_.* But generally a number of different *F_k_*(*x*) and *C_k_*(*x,y*) functions may be conceived depending on the specific plant ecological case. Alternatively, if limited or no prior information of the selection forces exists then model (1) may be fitted using e.g. spline functions (also see Laughlin et al., 2015).

The effects of the *m* traits on plant population growth is here assumed to be additive; but see the later discussion on the possibilities of relaxing this important assumption.

The different selection models may be fitted to longitudinal plant relative abundance data by specifying the relevant likelihood function. Since the predicted cover of plant species *j* in eq. 1 is not bounded between zero and one, the predicted cover was fitted to the observed cover using a normal distribution, where the standard deviation was scaled by the observed cover times one minus the observed cover, i.e. *ε*~*N*(0,*q*_*j*_(1 − *q*_*j*_)*σ*) For example, in the case of fitting the directional selection, models (2a) and (3a) are inserted into (1) the resulting likelihood function is,

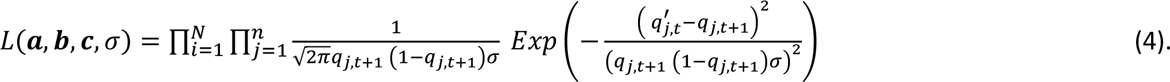

Most importantly, the parameter, *σ*, estimates the structural variance in the change in cover, which is the variance that is not explained by the model (1), and may consequently be used to get an estimate on the quantitative importance of the simplifying assumptions used in the modelling process.

## Demonstration with a Toy Example

In order to present and discuss the nature of the underlying assumptions and illustrate the possible use of the model, the above-outlined method was applied on a simple toy example of a realistic data type.

An arbitrary species-traits matrix with ten species and three traits, ***T***_10,3_, was constructed with random integer values (Table 1) and using an arbitrary directional selection scheme of the population growth based on the values of the three traits, 5(x_1_ − x̅_1_) + 3(x_2_ — x̅_2_) − 2(x_3_ − x̅_3_) + 100, where x*_k_* is the trait value of trait *k*. The initial cover of ten species was generated for a hundred plots using a Dirichlet distribution with all parameters set to one. The selection scheme was used on the generated initial covers of the hundred plots and afterwards normalized to obtain resulting cover values that sum up to one for each plot. Only the species-trait matrix, ***T***_10,3_, and the initial cover data, ***q***_*i*,1_, and resulting cover data, ***q***_*i*,2_, at plot *i* (*i* = 1,…100) were used in the further analysis, thus resembling the conditions in a real plant ecological study.

**Table 1.** The used species-traits matrix with ten species and three traits, ***T**_10,3_.*

| Species | Trait 1 | Trait 2 | Trait 3 |
| --- | --- | --- | --- |
| 1 | 3 | 2 | 3 |
| 2 | 2 | 4 | 2 |
| 3 | 4 | 1 | 6 |
| 4 | 7 | 5 | 3 |
| 5 | 2 | 3 | 8 |
| 6 | 9 | 4 | 4 |
| 7 | 2 | 3 | 6 |
| 8 | 4 | 6 | 5 |
| 9 | 6 | 3 | 1 |
| 10 | 5 | 1 | 5 |

The joint posterior distribution of the parameters in likelihood function (4) was simulated using a Bayesian MCMC algorithm (Metropolis-Hastings), where the parameters were assumed to have a uniform prior distribution, except for *σ,* where the prior was assumed to have an inverse gamma distributed with the parameters 0.001 and 0.001. The MCMC iterations had fair mixing properties and were judged to have converged to a stable joint posterior distribution after a lag phase of 50.000 iterations (results not shown). The joint posterior distribution was estimated from 50.000 iterations after the lag phase.

Statistical inferences on the individual parameters were based on the 95% credible intervals of the marginal posterior distributions. All calculations were done using *Mathematica* version 10 (Wolfram, 2015).

The generated cover data was successfully fitted by likelihood function (4) and the marginal posterior distributions of the parameters are summarized in Table 2. There were significant differences among several of the growth parameters and all nine growth parameters differed significantly from zero (Table 1). This indicates, although by using artificially generated plant cover data, that it is possible to estimate the effect of traits on population growth with an acceptable signal-to-noise relationship when fitted to hundred plots, which is a realistic number of replicates in ecological studies.

**Table 2.** The marginal distribution of the parameters of likelihood function (4) summarized by their 2.5%, 50%, 97.5% percentiles and the probability that the parameter is larger than zero.

| Parameter | 2.5% | 50% | 97.5% | P(X > 0) |
| --- | --- | --- | --- | --- |
| $a_1$ | 0.0911 | 0.0952 | 0.1011 | 1 |
| $a_2$ | 0.0907 | 0.0957 | 0.1003 | 1 |
| $a_3$ | 0.0615 | 0.0658 | 0.0696 | 1 |
| $b_1$ | 0.0019 | 0.0038 | 0.0063 | 1 |
| $b_2$ | 0.0001 | 0.0012 | 0.0032 | 0.986 |
| $b_3$ | 0.0033 | 0.0052 | 0.0070 | 1 |
| $c_1$ | -0.0894 | -0.0817 | -0.0740 | 0 |
| $c_2$ | -0.2092 | -0.1968 | -0.1818 | 0 |
| $c_3$ | -0.3053 | -0.2933 | -0.2820 | 0 |
| $\sigma$ | 0.0941 | 0.0984 | 0.1029 | 1 |

The covariance matrix of the joint posterior distribution and the graphs of the parameter iterations (not shown) showed almost no covariance between *a*_*k*_ and *c*_*k*_. This generally indicates that it is possible to partition direct population growth and population growth that is mediated by interspecific interaction.

## Discussion

Most importantly, a number of quite specific assumptions on the nature of selection and how the different traits interact (eqn. 1, 2 and 3), is needed to set up the model and to meet the objectives of the empirical modelling, i.e. to estimate the selection forces on traits while at the same time to discriminate between environmental filtering and niche partitioning processes. Such a modelling approach is in sharp contrast to the more simple and elegant maximum entropy models, where it is not necessary to specify detailed models on selection forces and how the different traits interact (Shipley, 2010a; Shipley, 2010b). Consequently, in the modelling approach presented in this study it is critical to assess or test the different necessary assumptions using either prior knowledge or model selection techniques.

As an additional tool in the model selection process valuable information may be obtained by estimates the structural variance, which is the variance that is not explained by the model and the underlying assumptions. If the structural variance is relative small then this is indirect evidence that the underlying assumptions to a certain degree are supported by the data. In the presented simple case-study the median estimate of the structural standard deviation was 0.0984 (Table 2), which should be compared with the expected cumulative cover changes of ten species with three traits. However, more worked-out empirical examples of real data are needed in order to assess the importance of this level of structural variation. Finally, the conclusions of the model should of course be compared with independent information or hypotheses on the nature of trait selection.

For simplification it is assumed in model (1) that there is no significant intra-specific trait variation (but see Laughlin et al., 2012) or intra-specific variation in population growth rate. Generally, using model selection techniques, it will be possible to test what type of selection (directional selection, stabilizing selection, or disruptive selection) is best supported by the data and, consequently, to generate and test hypothesis on trait based assembly rules and possible mechanisms underlying plant species coexistence. Furthermore, if plant abundance of perennial plants is measured several times during a growth season, e.g. in spring and autumn, then the trait selection processes during summer growth may be estimated independently from the trait selection processes during over-wintering and, consequently, allows the generation and testing of temporal coexistence mechanisms (storage effects, Chesson, 2000).

Regarding the used assumption on the interactions between traits, model (1) assumes additivity among the traits in regulating population growth. Generally, little information exists on the interaction among traits (Kraft et al., 2015b), but the covariance matrix of the estimated selection coefficients *a_k_* and *c_k_* may give important insight on the selection forces operating on a suite of correlated plant traits, e.g. specific leaf area and leaf dry matter content, as previously demonstrated by e. g. Lande and Arnold Lande (1983). Again, the above-discussed model selection techniques may be used to discriminate between different hypotheses, and in the case that some modes of interactions are not supported by data it may be concluded that new ecological insight has been established.

In the presented simple demonstration case, the used cover data were generated assuming a uniform environment, but if the cover data had been sampled along an environmental gradient, then the selection models (2) can be made dependent on the environmental gradient; and the effect of traits on population growth can then be estimated as functions of the environmental gradient. In similar ways, the selection models (2) can be modified to fit many different ecological circumstances and the demonstrated model in this paper is only one possibility of a large class of models that may be fitted using the outlined methodology. The model is currently being used to examine the effect of plant competition on trait selection along a hydrological gradient (Damgaard et al, in prep.)

Generally, it will be possible to generate ecological predictions with a known degree of uncertainty from the outlined trait selection model by inserting values from the joint posterior distribution of the parameters into numerical iterations or a numerical solution of equation (1). Such ecological predictions may be used directly in applied plant ecological questions, e.g. effects of climate change, pesticides, or nitrogen deposition on plant communities.

The outlined trait selection model is a one-step trait selection process that only operates locally and is, thus, conceptually simpler than the two-step process meta-community model that previously has been used (e.g. Bernard-Verdier et al., 2012; Webb et al., 2010). One of the advantages of this simpler model is that it allows ecological predictions to be generated without knowledge on meta-community dynamics which, typically, is unknown. The data requirements of the presented model are modest, i.e. time series data on plant species abundance and a species-trait matrix. Consequently, the model may be used to model trait selection, including the effect of interspecific interactions, in many existing plant ecological datasets. Naturally, the method is extendable so that time series longer than two years or time series data with irregular sampling intervals also may be fitted.

In the used modelling approach interspecific interactions are measured directly using time series plant abundance data as the effect neighboring plants have on growth (Damgaard, 2011; Damgaard et al., 2009; Damgaard et al., 2013; Damgaard et al., 2014), and this allows us to model the underlying ecological processes. In my opinion, the filter analogy has been overused in empirical plant ecological trait literature, e.g. when loosely referring to a “competitive filter” or “biotic filter” without specifying the details of the underlying ecological processes (Kraft et al., 2015a). Since multiple assembly processes can lead to the same pattern of trait dispersion and the same process can lead to different patterns of trait dispersion (Herben and Goldberg, 2014), it is a clear advantage of the outlined model that it operates on the process level and that it is possible to mathematically describe the details of different ecological processes within the framework.

## Acknowledgement

Thanks to Zdeněk Janovský and anonymous reviewers for valuable comments on a previous version of the manuscript.

